# Latitudinal phylogenetic and diversity gradients are explained by a tropical-temperate transitional region

**DOI:** 10.1101/2025.02.23.639794

**Authors:** David Lerner, Jason T. Weir, Tamir Klein, Gili Greenbaum

## Abstract

The distribution of ecological and evolutionary forces throughout space bring about the patterning of biodiversity. In large geographical areas, this causes the regionalization of biodiversity into structured units known as bioregions. In order to understand how such patterns emerge, a clear delineation of bioregions is required. We use tree species as model taxa in order to analyze the global distribution of biodiversity and understand how latitudinal gradients of biodiversity, specifically the latitudinal phylogenetic and diveristy gradients are formed. By compiling an extensive dataset of tree species distributions and their phylogenetic relationships, we use a data-driven approach to delineate global bioregions of similar evolutionary histories, termed phyloregions. Our analysis reveals the presence of a region between the tropical and temperate regions, coined ‘bridge’ phyloregion, with a unique evolutionary composition and characteristically weaker association to climatic and environmental parameters. Through simulations, we show that the presence of latitudinal phylogenetic and diversity gradients are much more likely to emerge in the presence of an independent ecological region between tropical and temperate regions, suggesting that its role as a stepping-stone in colonization of species between distinct climatic zones has shaped latitudinal gradients. This study highlights that accurate delineation of evolutionary structures of biodiversity can reveal previously cryptic regions with fundamental evolutionary roles in the formation of biodiveristy patterns.

**Significance Statement:** Biodiversity patterns are shaped by geography, forming distinctive units known as bioregions. At global scales there are important gradients in terms of species diversity and phylogenetic relations that are well documented, but the underlying processes that generate them are unclear. We comprised an extensive dataset of tree species distributions and phylogeny, and identified the bioregions from the dataset. We analyzed the diversity and the phylogenetic gradients and identified a previously cryptic bioregion between tropical and temperate zones. Though less climatically distinct, we show through simulations that the presence of this transition zone better explains latitudinal biodiversity patterns. Our findings refine the global dispersal dynamics of species between tropical and temperate regions and highlight the role of this intermediate region in shaping global biodiversity patterns.

## 2 Introduction

Biogeographic patterns emerge primarily out of the interaction between ecological and evolu-tionary forces [1, 2, 3, 4, 5]. Efforts to map diversity patterns and to delineate and characterize the formation of bioregions have been ongoing for over a century ([6, 7, 8]), but the recent availability of global climatic data, large phylogenies, and detailed occurrence data now makes it possible to uncover and investigate biogeographical patterns in detail [9, 10, 11]. One of the key axes along which biogeographic patterns emerge is latitude, with an important example being the diversity gradient. However, the underlying ecological and evolutionary processes that have led to the emergence of many latitudinal gradients remains unclear.

Many types of latitudinal gradients have been described, including gradients in body size [12, 13], genetic diversity [14, 15], and avian colourfulness [16]. While some latitudinal gradients have been well-characterized and attributed to environmental factors (e.g., larger body size in colder climates, higher primary productivity in the tropics due to higher solar radiation and rainfall), other gradients remain harder to explain. This is the case for two of the main broad and fundamental latitudinal gradients: the latitude diversity gradient [17, 18], where lower latitudes are characterized by higher species and genera diversities, and the latitudinal phylogenetic clustering gradient [19, 20], which is usually interpreted in terms of phylogenetic niche conservatism, where lower latitudes are characterized by higher phylogenetic clustering [21, 22, 23].

Although substantial differences in levels of phylogenetic clustering between regions have been observed in many taxonomic groups [24, 20, 25], the underlying mechanism for this pattern is still unresolved. One explanation for such biogeographic patterns suggests that environmental conditions in tropical regions are conducive of niche conservatism, where new species emerging in the tropics tend to be adapted to the high precipitation and temporally ho-mogeneous environment of their ancestors, more than in temperate regions. [23, 24, 26]. This tropical niche conservatism results in more limited colonization capacity of tropical species [27, 21], thus generating phylogenetic clusters with over-representation of tropical species. Yet, other suggested mechanisms that do not include niche retention have also been suggested, in-cluding region-related evolutionary rates [28, 29, 30], species antagonism and competition [31, 32], or niche shifting [30, 33] (see [21] for a comprehensive review). Importantly, it has been suggested that the latitudinal diversity gradient may itself be a consequence of the latitudinal phylogenetic gradient, because higher phylogenetic clustering in the tropics may mean that the tropics act both as ‘cradles’ of species, (i.e., higher speciation rates), and ‘museums’ (i.e., lower extinction or colonization of tropical species) [34, 20, 35, 36]. In order to clarify the underlying mechanism of the latitudinal phylogenetic gradient, and its relation to the diversity gradient, it is important to consider how the interactions between different evolutionary forces may generate these global patterns [22, 37, 21], by integrating phylogenetic data into biogeographical studies.

A main challenge in understanding biogeographical patterns is that delineating regions is usually a prerequisite for any comparison between geographic regions, and region delineation often requires *a priori* assumptions. In most cases, gradients have been studied using either simplistic delineation systems (e.g. ‘tropical-temperate’ dual delineation; [20], or by partitioning the world into predetermined ecoregions; [38, 37]). However, *a priori* delineations of regions may overlook important biotic regions and their eco-evolutionary effects on biodiversity gradients. One potential way to overcome this limitation is to utilize data-driven approaches for region delineation [39, 40, 41]. Such methodologies reduce the need for assuming we know what the important bioregions are. It may also ensure that no cryptic bioregions, which may be important for understanding how latitudinal gradients are formed, are missed. In addition, when studying the latitudinal phylogentic gradient, it is important to incorporate phylogenetic distances in the region delineation process because species are not phylogenetically independent[42, 43], and treating them as such may generate artifacts. Therefore, generation of ‘phyloregions’ using data-driven approaches that incorporate phylogenetic distances may correct for phylogenetic structure when determining bioregion classification.

Trees are one of the most widely used taxonomic groups for studying biogeographical patterns, for several reasons. Firstly, tree species are widely distributed, occurring in most terrestrial regions and are more easily sampled compared to less sessile taxa, and are, therefore, ideal for characterizing global patterns. For similar reasons, genomic data is readily available for most tree genera, and therefore large and reliable phylogenies can be constructed for most tree groups. By integrating distribution and phylogenetic data, several studies have investigated the processes that generate biodiversity patterns at large regional and global scales ([26, 9, 44], highlighting the observational patterns of tropical phylogenetic clustering. Understanding tree biogeography is of particular importance because many foundational global biodiversity classifications, such as the biome system [8, 45], rely heavily on vegetation patterns, and in particular tree distributions [46, 47]. Finally, the unique evolutionary history of tree species generated clearly defined biodiversity structures, such as the latitudinal distinction of gymnosperms almost exclusively in high latitudes and angiosperms dominating the low latitudes due to competitive clade replacement [48, 49].

Here, in order to understand how evolutionary forces shape biodiversity patterns, we complied and analyzed an extensive dataset of tree species from across the globe. We apply a data-driven approach to delineate and characterize the main phyloregions of trees. Using this characterization, we study the latitudinal phylogenetic gradient, and identify intermediate ‘bridge’ regions between ecologically well-defined tropical and temperate regions. To better understand the role of the ‘bridge’ region in the formation of diversity patterns, we design a modeling approach to study how macro-evolutionary forces can give rise to the specific characteristics of the latitudinal gradients we observe.

## 3 Results

### 3.1 Global phyloregions of tree species

In order to identify worldwide phyloregions for tree genera, we compiled an extensive dataset of geographic distributions of 2582 species (obtained from [50]), as well as phylogenetic relationships resolved at the genus level (obtained from [41]), result in a final dataset of 481 genera for which geographic and phylogentic data was available. We then used an unsupervised hierarchical clustering method aimed at minimizing phylogenetic *β*-diversity distances, based on the genera composition from independent geographical units. This method effectively delineates regions characterized by similar evolutionary histories and phylogenetic relationships, resulting in a map where each cluster represents a distinct phyloregion. To determine the number of regions, we used the elbow criterion, which suggested *K* = 9 clusters. We observe that, although phyloregion assignment was made for each geographic location independently, the inferred phyloregions are composed of large continuous geographic regions, and that most phyloregions are present in different continents (Fig. 1a). The tropical regions (especially in the New World) were represented by a single phyloregion (cluster 9 in Fig. 1a), while some of the higher latitude phyloregions (clusters 5, 7 and 8 in Fig. 1a) were identified in both northern and southern hemispheres. This pattern reflects a relatively symmetrical global distribution of phyloregions with respect to the equator. The same pattern was observed for a lower number of clusters (*K* = 6) (Fig. S1).

**Figure 1:**
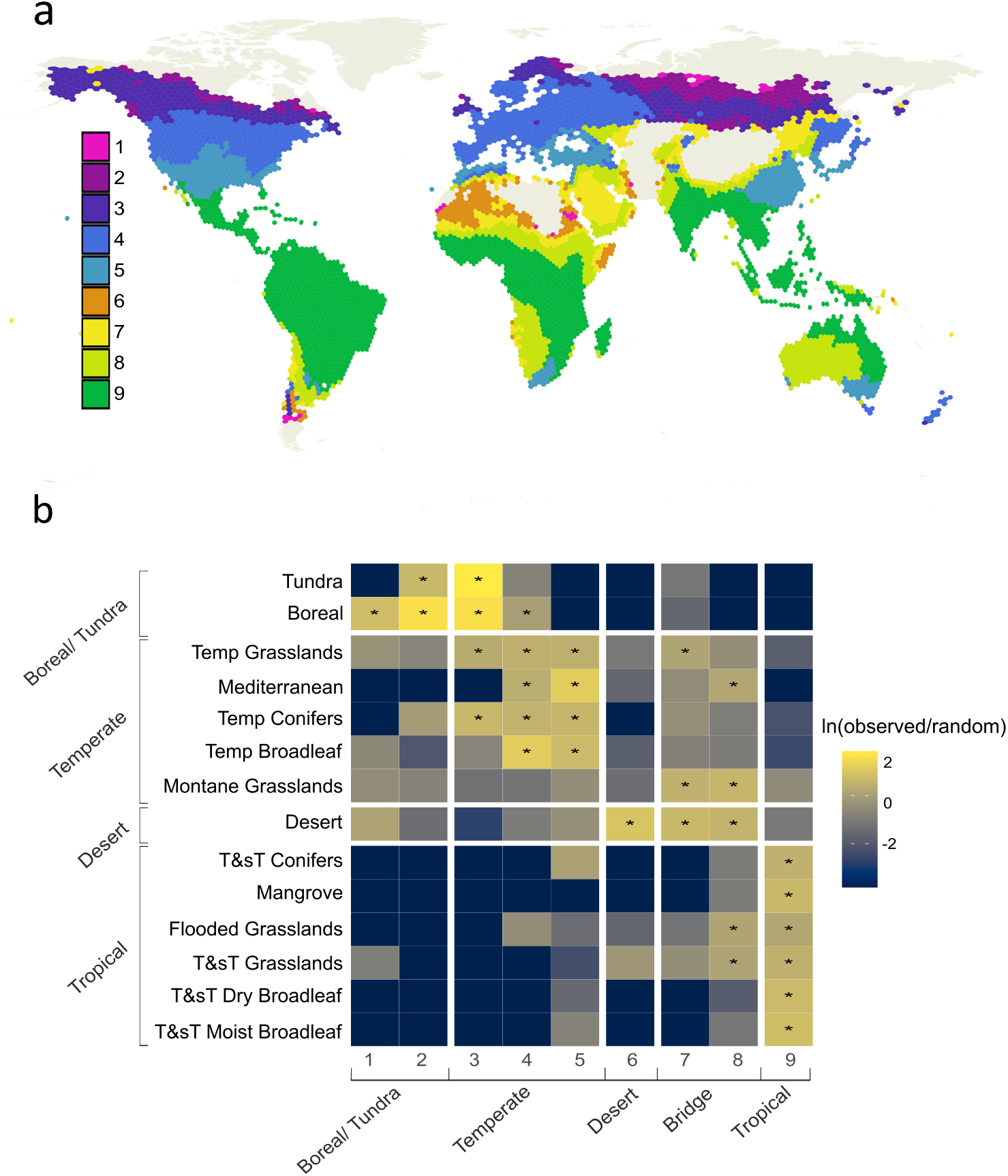
Phyloregion delineation of tree species. (a) Nine inferred phyloregions of tree species. The phyloregions are inferred using a clustering analysis of 5851 geographical locations, taking into account the genera compositions and phylogenetic distances. (b) Proportion of overlap between each ecological biome and the nine phyloregions, compared to a randomized phyloregion bootstrap. The heat map shows the natural logarithm of the ratio between the actual overlap and the bootstrap overlap, with yellower colors signifying higher-than-random overlap. Statistical significance of high overlap is marked with an asterisk (*p <* 0.01). Based on the overlap with biomes, we assigned the nine clusters to five phyloregion types, denoted below the heatmap.

We characterized the phyloregions in terms of ecological features by comparing the geo-graphical distribution of phyloregions with that of worldwide biomes (Fig. 1b). Each phylore-gion showed significant geographic overlap with at least one biome, and we were, therefore, able to denote the ecological span of each phyloregion. Some clusters were associated with specific biomes: cluster 9 was exclusively associated with tropical and subtropical biomes, cluster 6 uniquely overlapped with the desert biome, clusters 3–5 were primarily associated with the temperate biomes, and clusters 1 and 2 were clearly associated with boreal-tundra biomes. In contrast, clusters 7 and 8 were not associated with any specific biome type, but rather spanned across multiple biomes, including tropical, temperate and desert. These two unique clusters, which we term ‘bridge phyloregions’, extend across grassland, desert and mountainous regions (Fig. 1b), spanning the Saharan, Arabian, Kalahari, central Asian and Atakama deserts, as well as mountainous regions of the Himalayas, the southern Andes and east Africa. The presence of bridge phyloregions as a transition between the tropical (cluster 9) and temperate (clusters 4 and 5) was observed throughout the old world and in the southern hemisphere of the new world. In the northern hemisphere in the New World, the bridge phyloregion is mostly absent, and the tropical phyloregion directly transitions to temperate except for the minor presence of cluster 8 in Baja California.

We repeated the analysis while including only geographic cells containing more than 30 genera, because small sample sizes may bias phylogenetic analyses [51]. The resulting phyloregion pattern is similar to the one obtained with the full dataset (Fig. 1), with clear presence of bridge phyloregions (clusters 2 and 3 in this analysis) that span tropical, temperate and desert regions. Because most of the low sample size cells that were excluded are in boreal, tundra, or desert regions, we do not observe phyloregions that specifically represent these biomes.

In terms of phylum classification, the distribution of taxa among phyloregions differs between tropical and temperate regions (Fig. S3a–b), with all temperate and boreal/tundra phyloregions (with the exception of cluster 5) composed primarily of gymnosperms, and a higher proportion of angiosperms in the tropical, bridge, and desert phyloregions. This last result is expected because gymnosperms generally predominate at higher latitudes, and angiosperms predominate in tropical, subtropical, and lower-latitude regions [52, 49, 53].

To study the evolutionary relationship between phyloregions, we measured the community and phylogenetic distances between the clusters using taxa *β*-diversity and phylo-*β* diversity metrics (Fig. 2c and Fig. S4). The phylogenetic and taxa turnover between phyloregions indicated a geographic-dependent distance between phyloregions, with the low-latitude tropical phyloregions at one extremity of the main axis and the high latitude tundra phyloregions at the opposing extremity (the exception are the desert phyloregion, cluster 6, and the high latitude boreal phyloregions, clusters 1 and 2). The sequential distribution of phyloregions along the first principal coordinate suggests a step-wise evolutionary divergence, where each phyloregion has the closest phylogenetic distance to its immediate geographic neighbor (although high-latitude clusters are located at both sides of the equator).

**Figure 2:**
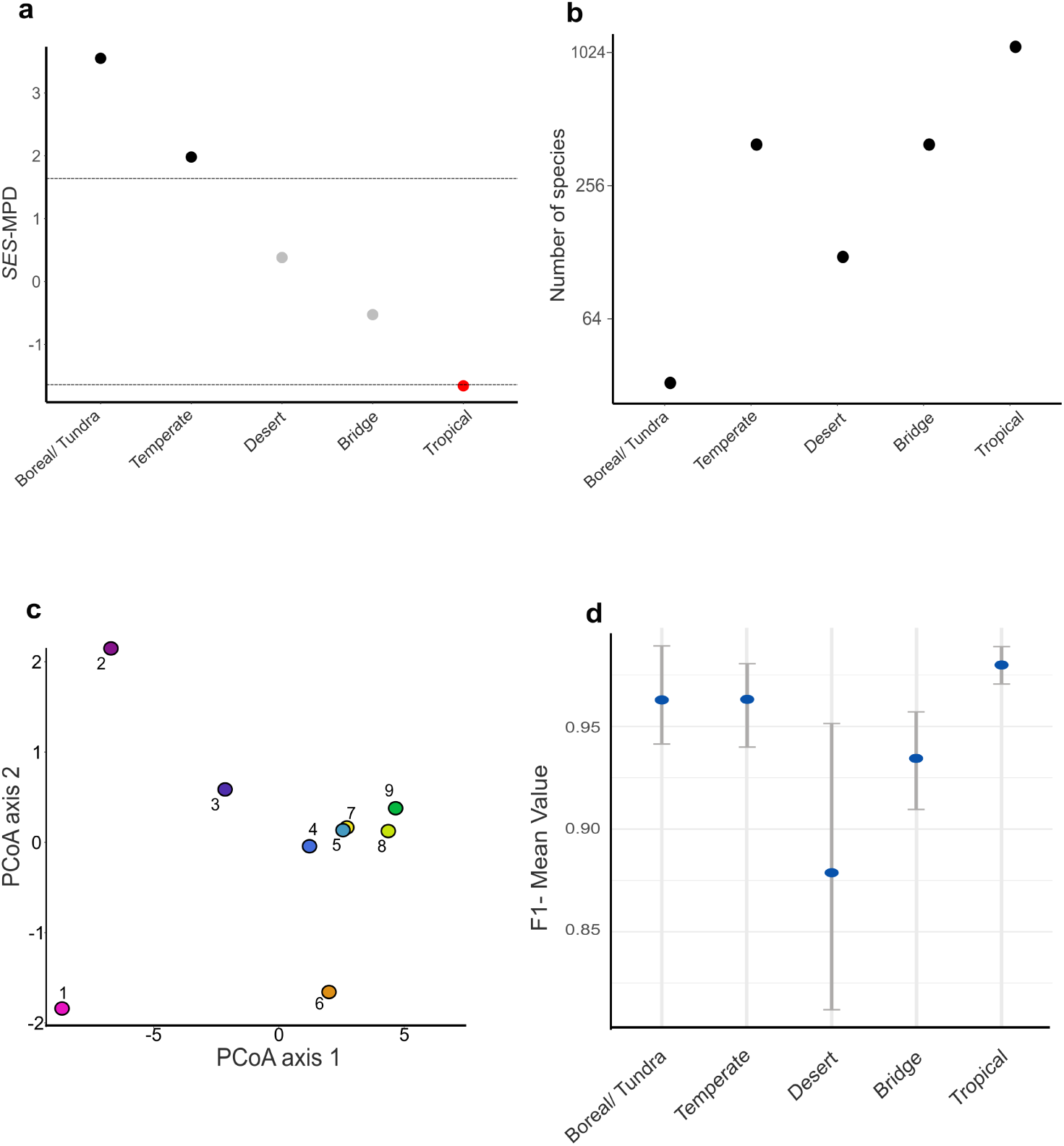
Evolutionary and ecological characteristics of tree phylogerions. (a) Phylogenetic diversity throughout the different phyloregion types, measured as SES-MPD Z-scores, for the five main phyloregion types identified. Phyloregions with SES-MPD *z*-scores *< −*1.96 are denoted with red dots, and represent significant phylogenetic clustering of tree species in these phyloregions. Phyloregions with SES-MPD *z*-scores *>* 1.96 are denoted with black dots, and represent significant phylogenetic over-dispersal of tree species. The values shown are the average value for the SES-MPD of all phyloregions assigned to a phyloregion type. (b) Genera diversity of phyloregions, measured as the average number of genera in each of the phyloregions types. (c) Evolutionary relationship among phyloregions in terms of Phylogenetic *β* diversity. Shown is a PCoA ordination plot of the phylogenetic distances between phyloregions, measured using Rao’s *D* index. (d) Relationship between environmental factors and phyloregions. A random-forest model trained with environmental variables was used to predict phyloregion type assignment, to determine the extent that phyloregions are associated with environmental conditions. Show are the means and 95% confidence intervals of the fraction of correctly identified phyloregion-type (F1-score), representing the ability of environmental data to predict phyloregion assignment.

### 3.2 The latitudinal diversity and phylogenetic gradients in relation to tropical niche conservatism

We quantified diversity of the main phyloregion groupings (tropic, desert, temperate, and boreal/tundra phyloregion groups) by genera counts, and phylogenetic clustering by taking the standardized effect size of the pairwise mean phylogenetic distance (SES-MPD) among the genera assigned to each cluster (hereafter referred to as MPD, for brevity). We observed both the latitudinal phylogenetic (Fig. 2a) and the latitudinal diversity (Fig. 2b) gradients. Genera diversity increased with increasing latitude, but there was no substantial difference in the number of genera between temperate and bridge phyloregions, and the desert region showed less diversity than expected from its latitude. For phylogenetic clustering, we observe a latitudinal gradient in MPD, marked by significant clustering in the tropical phyloregions, transitioning to an over-dispersed phylogenetic arrangement in the boreal/tundra phylore-gions (Fig. 2b). This pattern was also observed when analyzing MPD at the biome level (Supplementary Fig. S5). We tested to see if the strong differential latitudinal patterning of angiosperms vs. gymnosperms —– specifically the higher presence of gymnosperms at higher latitudes (Supplementary Fig. S3b) — affected the emergence of the latitude phylogenetic gradient. When gymnosperms were excluded from the analysis, using all geographic cells, the over-dispersion at higher latitudes, especially in the boreal/tundra phyloregions, was lower and even significantly clustered (Fig. S6), and only the desert phyloregion was not significantly clustered. However, when using only cells with *>* 30 genera, which should be less biased due to low sample sizes in some cells, the latitudinal gradient was similar to that observed when including both gymnosperms and angiosperms (Fig. S2). This suggests that, although the distributions of gymnosperms play a role in shaping the latitude phylogenetic gradient, the gradient is still present regardless of the latitudinal distribution of tree phyla.

Next we wanted to evaluate the extent of niche conservatism of phyloregions, without considering phylogenetic clustering. One perspective through which this can be done is by evaluating how strongly the environment dictates the distribution of phyloregions, because a strong environmental influence would suggest that closely related species are retained within a given environmental niche. Conversely, if environmental factors are less influential, it would imply that new species or genera are not constrained to remain in an environmental niche of their ancestor. To quantify the extent to which environment explains distributions, we con-ducted a random-forest classification analysis using present-day environmental factors (e.g., climate, elevation, soil-available phosphorous), aimed at predicting genera compositions for geographic locations from environmental data. In general, we found that environmental fac-tors are strongly predictive of phyloregion assignment for geographic locations (accuracy of model = 91%). However, prediction accuracy varied substantially for different phyloregions, with the model accurately predicting the distribution for the tropical, temperate and boreal phyloregions, but less so for the bridge and desert phyloregions (Fig. 2d). A similar random classifier model, treating all phyloregions as independent clusters, as opposed to the main phy-loregion groupings (boreal/tundra, temperate, desert, bridge and tropical), produced similar results for all except for phyloregion 1 (Fig. S8). This low predictive accuracy of phyloregion 1 could suggest that this phyloregion, although primarily composed of boreal regions, also includes regions that were not successfully classified into any other phyloregion (Fig. 1a), mainly within deserts (miss-classification of regions that do not fit into any other category is a common limitation of hierarchical clustering techniques [54]).

Taken together, these results are consistent with a atitudinal phylogenetic gradient (Fig. 2a), but we do not find evidence that niche conservatism is its driving mechanism. We observe that the bridge phyloregion is characterized as a transition zone between temperate and tropical areas (Fig. 2a) in terms of phylogenetic clustering, geography and diversity, but that it spans across several ecological biomes and is less strongly determined by environmental conditions than the other phyloregions (Fig. 2d). These features challenges our understanding of how the latitudinal phylogenetic gradient is formed.

#### 3.2.1 The role of the bridge region in the emergence of latitudinal gradients

To better understand how the attributes of the transitional bridge phyloregion impact the macro-evolutionary processes driving the latitudinal phylogenetic and diversity gradients, we designed a modeling framework that incorporated key macro-evolutionary forces — speciation, extinction, and colonization. We model three regions, representing the tropics, the temperate and the bridge, where speciation and extinction occur at given rates in each region, but colonization can occur only between the bridge regions and the other two regions. For comparison, we also modeled the same scenario without the bridge region, with only two regions representing the tropics and the temperate between which colonization can occur. We simulated a birth-death speciation-extinction process with colonization, and noted the MPD values for each region at the end of each simulation, defined as the time for which a total of at least 500 extant species was reached.

Because climatic tolerance is an important regulator of diversification rates [55, 56, 57], we specified speciation and extinction rates to the tropical and temperate regions. To reflect the weak environmental association that we observed for the bridge phyloregion (Fig. 2d), we did not set diversification rates for this phyloregion, and instead genera dispersing to the bridge phyloregion kept the rates from their ancestral region. We used these simulations to understand how the inclusion of a transitional bridge region affected phylogenetic measures by identifying parameters that generate a gradient from over-dispersed to under-dispersed phylogenetic distances (Fig. 3a), as observed in Fig.2b, while maintaining a latitude diversity gradient. We compared simulations with and without a bridge region by analyzing the proportion of the simulations that resulted in two conditions: (1) a latitudinal diversity gradient, whereby there is a gradient of genera richness, from highest in the tropical region to lowest in the temperate region (denoted as ‘condition 1’); (2) the formation of a latitudinal phylogenetic gradient, defined by a high MPD difference between tropical and temperate regions (denoted as ‘condition 2’. This condition was defined to be satisfied when the MPD in the tropical region was in the bottom 20% out of all MPD values attained in our simulations, and the temperate MPD was in the top 20% of MPD values; we also evaluated a more stringent of this condition, denoted as ‘condition 2*’, where MPD values in the temperate and tropics were required to be in the extreme top or bottom 10% of values, respectively. For simulations that included the bridge region, we also required that *MPD_trop_ < MPD_bridge_ < MPD_temp_* to ensure a gradient of phylogenetic clustering (Fig. 3a).

**Figure 3:**
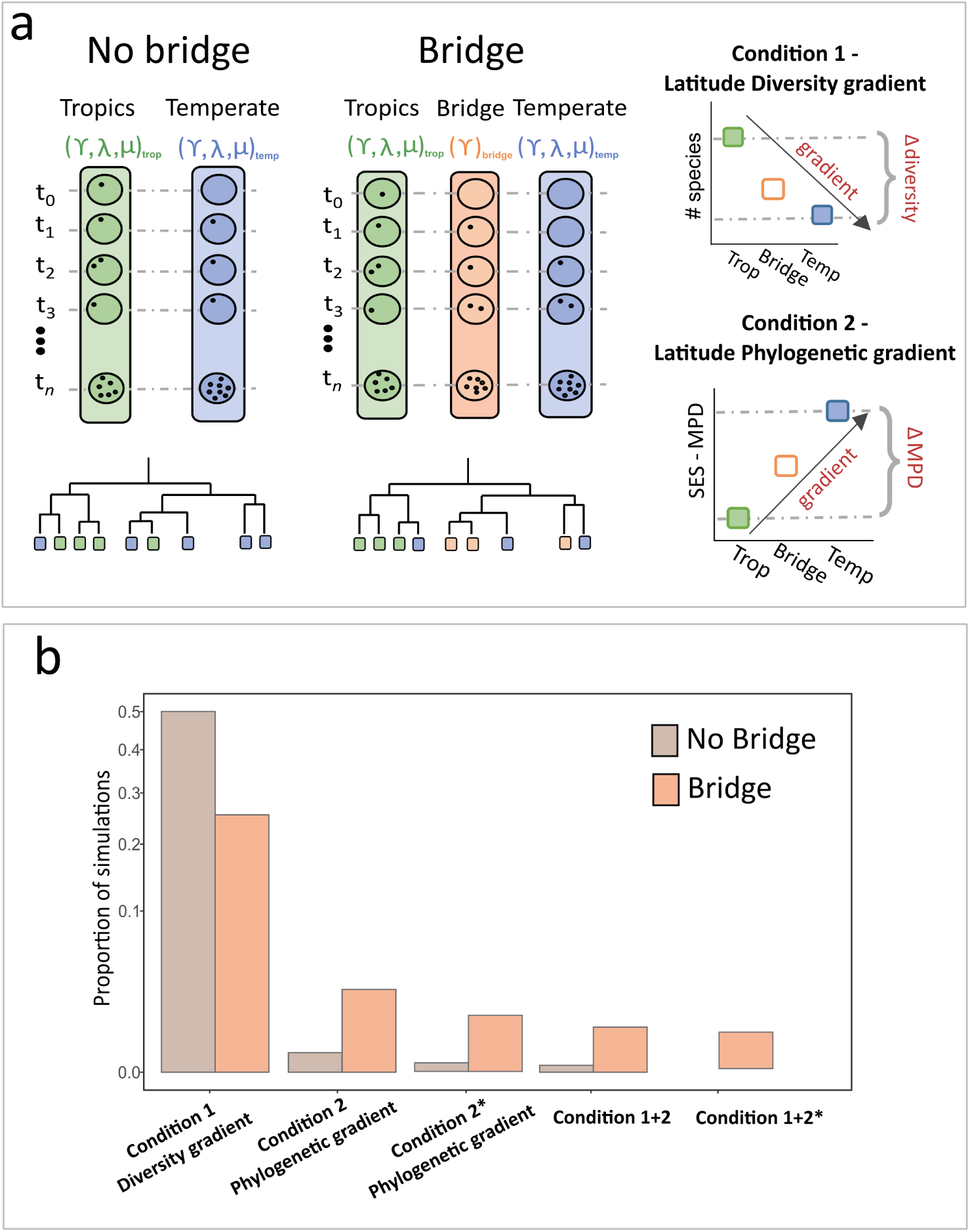
The role of the bridge phyloregion in a simulation model. (a) A schematic representation of our modeling framework for studying the role of the bridge phyloregion in generating latitudinal gradients. We model 2–3 regions, representing the tropical (green), temperate (blue), and bridge (orange) regions. In one set of simulations we include the bridge region, and in the other exclude it. For a set of parameters (speciation, extinction for each regions, and dispersal rates between regions) we track the phylogeny, and compute the standardized effect size mean phylogenetic distances (SES-MPD) of the regions, as well as genera diversity. We consider the outcomes of a (i) a latitude diversity gradient (condition 1), (ii) a latitude phylogenetic gradient (condition 2) and (iii) a combination of phylogenetic and diversity gradient (Condition 1 and 2). (b) The proportion of the simulations for which the latitudinal diversity gradient, phylogenetic gradient, and the combination of both emerged in the simulation including and excluding the bridge regions. Two versions of condition 2 were evaluated, where 2* is more stringent than condition 2. *y*-axis is scaled using a square root transformation to allow visualization of cases with a small proportion of simulations.

Although the model without a bridge region had a higher fraction of simulations that met condition 1 alone (only diversity gradient), the inclusion of the bridge region substantially increased the proportion of simulations that satisfied both conditions 1 and 2 (Fig. 3, Fig S10), with a 30-fold increase compared to the model without the bridge region. Under the more stringent condition 2*, none of the simulations that did not include the bridge region resulted in both conditions being met. This result highlights the role that the bridge region plays in generating phylogenetic patterns, and may indicate that patterns such as those presented in Fig.2b are unlikely without such a transitional region between tropical and temperate regions.

#### 3.2.2 Evolutionary dynamics leading to the latitude phylogenetic and diversity gradients

To further characterize the scenarios in which the latitudinal gradients emerge, we partitioned the parameter space into categories based on the colonization dynamics each configuration represents. For simulations including the bridge phyloregion, we define five colonization classes: (i) ‘equal colonization’, where all colonization rates are the same, (ii) ‘out-of-the-tropics’, where net colonization is from the tropics outwards, (iii) ‘into-the-tropics’, where net colonization is from the temperate region outwards, (iv) ‘out-of-the-bridge’, where colonization tends to occur from the bridge outwards towards the tropics and temperate regions, and (v) ‘into-the-bridge’, where colonization tends to occur from the tropics and temperate regions into the bridge (Fig. 4a shows the exact conditions used to partition the parameter space into the five categories). For the simulations that excluded the bridge phyloregion, we used only three categories: ‘equal colonization’ (*γ_tropic→ bridge_* = *γ_temperate→ bridge_*), ‘out-of-the-tropics’ (*γ_tropic → bridge_ > γ_temperate → bridge_*), and ‘into-the-tropics’ (*γ_tropic→ bridge_ < γ_temperate → bridge_*). We then compared the proportion of simulations for which each latitudinal gradient, diversity or phylogenetic, emerges, as well as the proportion of simulations in which both emerge.

**Figure 4:**
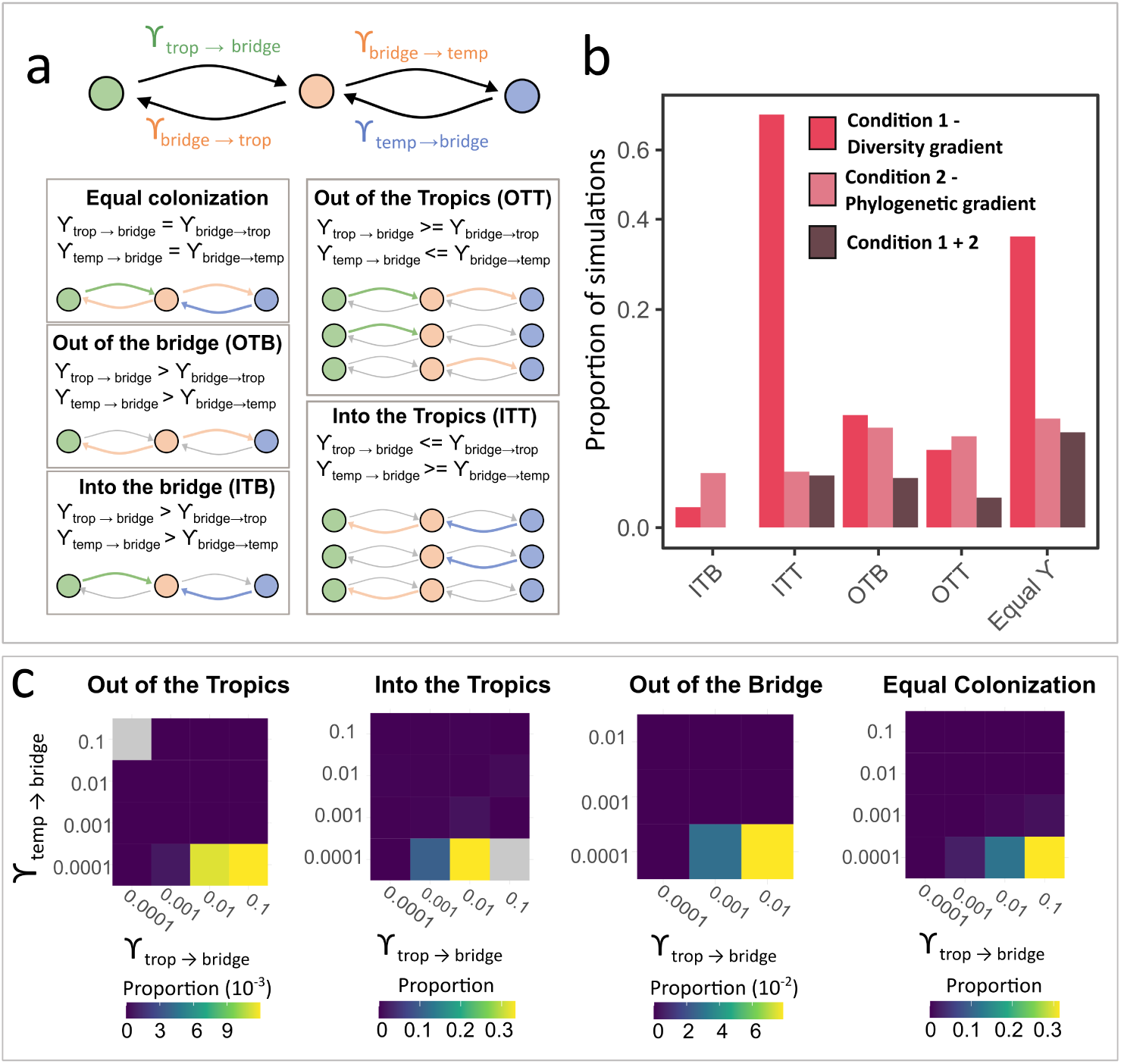
Scenarios in which latitudinal gradients emerge. (a) The parameter space for our simulations was partitioned into five categories based on colonization pattern configurations when the bridge region was included, and three categories (‘out-of-the-tropics’, ‘equal colonization rates’, ‘into-the-tropics’) when the bridge region was excluded. (b) The proportion of the simulations for which the latitude diversity gradient, the latitude phylogenetic gradient and a combination of both emerged, for each colonization category. *y*-axis is scaled using a square root transformation. (c) Analysis of the proportion of colonization configurations that generate both latitude gradients. Parameters *γtrop → bridge* and *γtemp → bridge* represent the colonization rates described in (a). The grey quadrants represent combinations that do not exist for this configuration of parameters under this colonization scenario.

Under four of the five colonization categories we observed simulations that met both latitudinal gradient conditions, whereas for the ‘into-the-bridge’ scenario no simulation resulted in both gradients (Fig. 4b). This result was robust to varying thresholds of extreme MPD values, obtained by comparing the highest and lowest proportions of MPD values between tropical and temperate regions (Fig. S9). Although ‘into the tropics’ scenario had the highest proportion of simulations producing a latitude diversity gradient, the ‘equal colonization’ scenario appeared as the scenario with the highest proportion of simulations that met both conditions. In contrast, the other three scenarios (‘out-of-the-tropics’,‘into-the-tropics’, and ‘out-of-the-bridge’), showed consistent differences in the proportion of simulations that resulted in both conditions (Fig. 4b, Supplementary Fig. S9). These results indicate that both latitudinal gradients emerge when the bridge region serves to connect the tropical and temperate regions, by acting either as a stepping-stone (equal colonization, ‘out-of-the-tropics’ and ‘into-the-tropics’) or as a source (‘out-of-the-bridge’) for species— but not as a sink (‘into-the-bridge’).

When looking at the combination of parameters that result in both the latitude diversity and phylogenetic gradients, we observed that there were higher proportion of simulations generating them for higher tropical than temperate speciation rates and higher colonization rates from tropical regions (*γ_trop→ bridge_*) than for temperate regions (*γ_temp → bridge_*), but no specific enrichment of extinction rate differences between tropical and temperate regions (Fig. 4c, Supplementary Fig. S10). An asymmetry in colonization rates was observed across all four colonization scenarios that generated both gradients, with greater tropical than temperate colonization (*γ_trop → bridge_ > γ_temp → bridge_*) (Fig. 4c). Surprisingly, this pattern emerged even under the ‘into-the-tropics’ scenario that was limited to higher temperate colonization rates (*γ_temp → bridge_ > γ_trop → bridge_*). This last result can occur when colonizations rates between tropical and bridge region (both *γ_trop→ bridge_* and *γ_bridge→ trop_*) are higher than those between bridge and temperate regions (both *γ_temp → bridge_* and *γ_bridge → temp_*) (Fig. 4a). Overall, our findings suggest that the emergence of both the latitudinal phylogenetic and diversity gradients is more likely to occur in the presence of a transitional niche-unconstrained bridge region, with higher tropical than temperate speciation and colonization rates.

## 4 Discussion

Using a data-driven approach, we identify five main distinct tree phyloregions, four corresponding to known ecological regions (tropical, desert, temperate, and boreal), and a fourth region at the geographic intersection between tropical and temperate regions (Fig. 1). This last phyloregion is characterized by intermediate phylogenetic clustering compared to the over-clustered tropical region and the under-clustered temperate regions, and is less associated with climatic and environmental parameters. We therefore term this region ‘bridge phyloregion’, since it seems to be a transition zone between two main eco-regions, rather than a clearly defined niche that determines evolutionary processes and rates. Our simplistic model demonstrates that the observed latitudinal diversity and phylogenetic gradients are much more likely to emerge in the presence of a bridge phyloregion that lacks phyloregion-dependent evolutionary rates. In addition, our simulations show that the bridge phyloregion most likely serves as a stepping-stone or source for species dispersal between tropical and temperate regions, rather than being a sink from species that transition to this region.

Considering the overlap of phyloregions and ecologically defined biomes, we observe that the bridge phyloregions is comprised of an area of land that spans tropical and temperate regions, with a strong presence in desert and grasslands (Fig. 1b, Supplementary Fig. S2). Notably, the bridge phyloregion was underrepresented in the desert and grasslands of North America (Fig. 1a), which raises the question of whether the bridge phyloregion is universal in all continent or arises in specific circumstances. One indication that the bridge region does occur in North America can be seen in our results with fewer clusters (Fig. S1) and more stringent filtering (Fig. S2), where the bridge phyloregion in North America is observed. Therefore, it may be that the bridge phyloregion signal is weaker in North America, and its emergence is more dependent on the paramaterization of the clustering analysis.

Although this is, to the best of our knowledge, the first study identifying a bridge phyloregion as an important transition area for species evolution and colonization between tropical and temperate regions, previous studies [58, 59] pointed out the importance of grasslands as stepping-stones in the colonization and further diversification of species between distinct climatic zones. The expansion of grasslands in the late Miocine and early Pleiocene transitional eras led to the increased colonization rates of tropical species [60, 61, 62]. Similarly, the inter-space between tropical and temperate regions was shown to harbor more colonization events to other regions than either the temperate or tropical regions [20]. These studies support our results of a transitional and diversification hub for species migrating between tropical and temperate regions. The result that the ‘into the bridge’ scenario is the only colonization dynamic that fails to produce both diversity and phylogenetic gradients (Fig. 4), suggests that the bridge phyloregion functions primarily as migration corridors for species moving between the tropical and temperate regions, rather that acting as sink habitats that retain colonizing species. Although the mechanisms by which bridge regions facilitate the colonization of both tropical and temperate species remain unclear, we suggest that the low-competition environments of grasslands and deserts and broad latitudinal spans of these regions act as buffer zones, facilitating dispersal, adaptation, speciation, and subsequent transitions to adjacent biomes.

### 4.1 The tension between the ‘tropical niche conservatism’ and ‘out of the tropics’ hypotheses

The ‘Out of the Tropics’ hypothesis [34], which postulates that the tropics act as ‘cradles’ for both tropical and temperate species, might seem mutually exclusive with the long-standing ‘tropical niche conservatism’ theory [63], which states that species that originate in the tropics tend to remain in the tropics. The two theories, therefore, can be viewed as assuming either high (’out of the tropics’) or low (’tropical niche conservatism’) colonization capabilities for tropical species, suggesting an apparent conflict between them. The tropical niche conservatism theory has been primarily supported by the latitudinal phylogenetic clustering gradient [22, 64, 35, 20], because the retention of species in the tropical niche should generate phylogenetic clustering in the tropics, relative to other regions. Our results reflect the tension between the two theories, because we show strong support for a latitudinal phylogenetic clustering gradient on the one hand (Fig. 2b), arguably strengthening the support for tropical niche conservatism, but our simulations show that higher tropical than temperate species colonization plays a key role in maintaining both the latitude phylogenetic and latitude diversity gradient (Fig. 4, Fig. 3, Supplementary Fig. S10).

One perspective in which the tension between the two theories can be resolved is by highlighting the following question — is phylogenetic niche conservatism a process or a pattern? In other words, is the retention of species within a specific niche the underlying evolutionary force leading to phylogenetic niche conservatism, or is it a phylogenetic signature that is the consequence of the interaction between other evolutionary forces [22, 21, 65, 66]? Ultimately, phylogenetic clustering is a tool by which we can quantify the similarity of traits and niches between closely related species [67, 68]. Our results show that both a latitudinal phylogenetic gradient and high colonization (emigration) of tropical species co-occur, which can be explained through the perspective that what is often referred to as phylogenetic niche conservatism could be merely a product of an underlying process, rather than the underlying process itself. Indeed, previous studies [28, 69, 70] have noted that phylogenetic signals leading to phylogenetic niche conservatism occur under evolutionary scenarios not unique to the decrease of species colonization rates, such as increased tropical extinction rates [28, 30]. Our results, therefore, support the idea that regions with labile species can still show a phy-logenetic clustering signature. Our model results suggest that observation of high clustering in the tropics do not require that tropical species are niche conserved, but rather that high colonization and adaptation capabilities to other regions of tropical species can explain the observed biogeographic patterns.

### 4.2 Diversification rates in relation to the phylogenetic and diversity lati-tudinal gradient

Previous studies have emphasized the significance of tropical niche conservatism and lower extinction rates in the tropics compared to temperate regions as the leading causes of the latitude diversity gradient [18, 19, 57]. However, a growing body of research is challenging the tropical niche conservatism hypothesis [28] and is highlighting a lack of consensus regarding lower tropical extinction as the primary driver of the latitudinal diversity gradient[71, 72, 73]. We observed that the latitude phylogenetic and latitude diversity gradients more often emerge under a combination of higher speciation rates in the tropics compared to temperate regions and directional colonization from tropical to temperate regions (Fig. 4, Supplementary Fig. S10). Therefore, our findings are in line with increasing evidence suggesting that the relationship between both speciation and extinction rates and latitude diversity gradients may be more complex than previously thought [71, 57].

A key limitation that arises from studying woody species exclusively rather than consider-ing the full range of plant taxa is that woody growth forms are not a monophyletic group, but instead, have independent evolutionary origins. This is due to the transitions between woody and herbaceous forms that have occurred over time [74, 11, 75]. These evolutionary reversals from woody to herbaceous forms may be misinterpreted as missing tips in the phylogeny, akin to extinction events. For example, the higher rate of these transitions at lower latitudes, al-though tested uniquely on insular woodiness [76], could reflect a potential bias in evolutionary rates in tropical regions [11]. However, the frequent occurrence of bidirectional evolutionary transitions between herbaceous and woody states across different taxonomic groups [75], as-suming they occur ubiquitously throughout the landscape, might counter this potential bias. While we, like others, have used woody species to study their evolution across biogeographical landscapes [20, 26, 9], further investigation is warranted to better understand the relationship between growth form transitions and evolutionary patterns.

Notably, our simulations managed to reproduce the latitude phylogenetic and diversity gradients even without explicitly incorporating the historical dynamics of gymnosperm clade displacement by angiosperms at low latitudes (Fig. 3). Despite the fact that we did not include complex mechanisms of clade turnover in our model, like sudden bursts of extinc-tion, we managed to reproduce the patterns of latitude phylogenetic gradient. Nevertheless, we do observe that clade displacement plays a role in the observed latitudinal phylogenetic gradient (Fig. 2 vs. Supplementary Fig. S6). However, previous studies have found that, in the New World, latitude phylogenetic gradient is observed in trees even when considering only angiosperms [20]. Interestingly, the evolutionary dynamics in the model that generate the latitudinal gradients we studied, higher tropical than temperate speciation coupled with tropical to temperate colonization, coincides with dynamics of the clade displacement of gymnosperms, whereby gymnosperms were excluded from the tropics by rapid tropical speciation of angiosperms and increased extinction of gymnosperms. Importantly, our modeling results suggest that the latitudinal phylogenetic gradient can emerge even in simplified models that do not account for the detailed evolutionary histories of whole clade speciation and extinction events.

## 5 Methods

### 5.1 Data set of tree species

This study was conducted by integrating the largest phylogeny to date of tree species [41], resolved to the genus level, including 787 genera, with a large dataset of tree species distributions [50], covering 3660 species from 1157 genera. The phylogeny of woody species was obtained from [41], whereby a tree was constructed using sequences from the *rbc*L and *mat* K plastid genes. This finalized phylogenetic tree, a collaboration from multiple previous efforts [26, 77, 78], is resolved to the genus level to prevent the misidentification of species [41, 9]. The distribution of tree species was obtained from [50], whereby polygons were generated from GBIF records, in order to represent the range of species. An integrated dataset, whereby each species was matched to their respective genus, generated a final phylogeny of 481 genera tips, including 2582 species of gymnosperms and angiosperms.

### 5.2 Generation of Phyloregions

A complete map of phyloregions was generated by computing the pairwise phylogenetic turnover given the species found at each global individual grid unit using a phylogenetic *β*-diversity metric. We initially divided the world map into grid cells using the ‘dggridR’ package, which employs the ISEA Discrete Global Grid System for equal-sized grid projection on a 2-dimensional plane. Each grid cell was approximately 23,000 km^2^.

Given the species composition within each grid, the pairwise phylogenetic turnover was calculated using Rao-H phylogenetic *β*-diversity [79, 51], a metric used to quantify the overall dissimilarity between two groups, in our case, grid cells, expressed as:

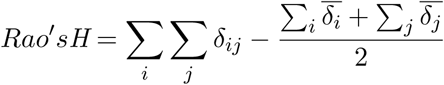

where *δ_ij_* is the most recent common ancestor (MRCA) between species in regions *i* and *j*. Species occurring in both communities have conspecific distances of zero, contributing to the mean pairwise turnover distance. We generated a 5722 x 5722 pairwise distance matrix. Due to the strong influence of species richness on diversity metrics [51], we excluded grid cells with fewer than five species to mitigate variance bias in species-poor regions such as deserts and high latitudes 1a), obtaining a final matrix that includes 4835 global grid cells. We used Rao’s *H* (derived from Rao’s *D*), as this metric has been shown to be highly correlated with other equally common metrices that calculate the phylogenetic distance within and between regions (e.g. MPD) [51].

Subsequently, we employed hierarchical clustering using the ‘complete’ method to minimize the maximum phylogenetic distance between each grid pair, thereby ensuring robust clustering based on phylogenetic similarities. To enhance the resolution of our clustering and reduce skewness in the data, we applied the natural log transformation to the outputs of Rao’s *H*. This transformation moderated the extreme differences within the distance matrix, facilitating a more detailed and fine-scaled clustering of phyloregions.

Using the ‘elbow’ method, we identified k=9 clusters in our hierarchical clustering analysis. This method involves plotting the total within-cluster sum of squares (WSS) against the number of clusters and identifying the point where the rate of decrease sharply shifts. This method allowed us to visually identify the most appropriate number of clusters, as it identifies the point at which adding more clusters does not significantly improve the variance explained by the model. The aim is to ensure that each phyloregion identified represents a distinct phylogenetic unit without over-fitting the model with an excessive number of clusters. In order to test for robustness, we reduced the choice of *k* clusters to 6, and observed very similar patterns (Supplementary Fig S1), as further noted in the Results section.

This method is described in detailed in [39].

All analyses were carried out in R (R Core Team, 2023), using the libraries ape [80], phytools [81] and picante [82].

### 5.3 Community ecology and statistical analyses

We included relative species abundance within each phyloregions in our analyses of phylo-genetc *α* and *β* diversity. Relative abundance was measured by comparing the observed versus expected species occupancy in each phyloregion.

Phylogenetic *β*-diversity between phyloregions was calculated in a similar manner as it was calculated to generate phyloregions (between grid cells), by creating an *n × n* pairwise matrix, where *n* is the number of phyloregions (9 phyloregions). Taxonomic *β*-diversity was also calculated between the phyloregions using the Sørensen Index.

The relative species abundance within each phyloregions was included in our analyses of phylogenetic *β* diversity between phyloregions. Relative abundance was measured by calcu-lating the relative presence of species occupancy in that phyloregion, given the number of grid cells within the phyloregion that the species occupies and the global presence of that species. Specifically, expected counts for each species within each phyloregion was estimated by dividing the observed species count in one phyloregion by the total counts of that species in all phyloregions, then multiplying the resulting proportion by the total species count in each phyloregion. A scaling factor was then applied by dividing each observed species count by the total count of that species, then multiplying the resulting proportion by the total species count in each region. This scaling factor was applied to ensure that the sum of expected counts matched the observed total counts for each region.

The *n×n* pairwise matrix distances for both *β*-diversity and phylo *β*-diversity were plotted on a 2-dimensional reduced representation, using either Non-metric Multidimensional Scaling (NMDS) for taxonomic *β*-diversity or Principal Coordinates Analysis (PCoA) for phylogenetic *β*-diversity, respectively. PCoA was chosen for phylogenetic analysis because it accommodates the actual distances calculated from the mean distance to the most recent common ancestor (MRCA) between phyloregions, which are critical for reflecting evolutionary relationships.

Phylogenetic *α* diversity was calculated using the mean phylogenetic distance (MPD) within phyloregions [67], much in the same way Rao’s D is generated:

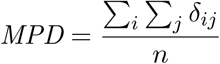

where *n* is the number of species within a community. In order to allow for a comparative analysis between phyloregions, we standardized the effect size of MPD values (SES)-MPD for each phyloregion, whereby:

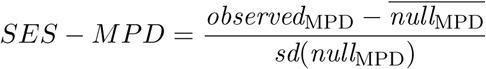

whereby *null_MPD_* is generated using a null model by randomizing the names of the taxa on the tips of the phylogeny. We chose such a constrained permutation technique as it generates a random phylogeny with the same phylogenetic architecture (branch length and distribution) as the actual phylogeny, thus reducing the Type I error rate that would be generated under a less constrained randomization technique, for example, by simulating novel phylogenies or randomizing the distribution of species within phyloregions. The permutation was carried out 500 times, in order to obtain the mean 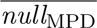 and standard deviation sd(*null*_MPD_) used for the model. A positive SES-MPD value represents higher phylogenetic distances than expected by chance, indicating phyloregenetic over-dispersal. Alternatively, negative SES-MPD values suggest shorter phylogenetic distances between the species compared to the null model, thereby implying phylogenetic clustering.

Next, to assess the influence of environmental niches on the distribution of phyloregions, we employed a Random Forest model. The model was set up to estimate how different environmental variables could predict the extent of phyloregions. We extracted the average value of each environmental variable in all grid cells. To ensure a balanced and representative training dataset, we randomly selected 80% of the grid cells from each phyloregion. This stratified sampling approach ensured that each phyloregion was equally represented in the training set. The remaining 20% of the grid cells from each phyloregion constituted the testing set. The environmental variables included were: Latitude, longitude, elevation, mean annual temperature (MAT), mean annual precipitation (MAP), temperature seasonality, isothermality and available phosphorous (Olsen P). All bioclimatic data was downloaded from WorlodClim Global Climate Data at a resolution of 5 arcminutes [83], extracted using the R package ‘raster’ [84]. Plant-available phosphorous was obtained from [85]. To validate the model and assess the reliability of its predictions, a bootstrap analysis was conducted. 100 bootstrap resampling of the original dataset were performed, with each sample drawn with replacement to match the size of the original dataset. For each bootstrap sample, the model was retrained, and its predictions were evaluated on a corresponding stratified test set, mimicking the original 80/20 split.

Model performance was evaluated using a confusion matrix for each bootstrap iteration. The confusion matrix provides a detailed breakdown of the model’s predictions, comparing predicted versus actual classifications. From the confusion matrices, we derived three key metrics: precision (the proportion of correct predicted identifications), recall (the proportion of positives that were correctly predicted, also known as sensitivity), and the F1-score (the harmonic mean of precision and recall). This process allowed the estimation of variability and confidence intervals.

### 5.4 Simulating the latitude phylogenetic gradient

#### 5.4.1 Model design

We study two model versions, one with two discrete ‘regions’ representing a tropical and a temperate region, and one with an additional ‘bridge’ region. Genera can occupy one or more regions at any time step, and can also move between adjacent regions (representing a colonization event). The three-region model is structured in a sequential arrangement, by allowing species colonization between tropical and temperate regions only by passing through the bridge region (Fig. 3a). In other words, colonization events in this model can only occur between the tropical and bridge regions, or between the temperate and bridge regions. Un-der this setup, we designed a spatially dependent non-zero sum extinction-speciation model, based on a previous study [86]. In our model, a genera originally occupies both the tropical and temperate regions. Speciation (*λ*), extinction (*µ*), and colonization (*γ*) are modeled as stochastic binomial processes, with region-dependent probability rates, whereby the chance of an event occurring is the per-lineage probability modeled at each time step, independent of the previous generation. In other words, at each time step each lineage has a fixed probability to speciate, a fixed probability to go extinct, and a fixed probability to colonize a different region. Therefore, regions with more genera will have a higher probability of speciation, extinction and outgoing colonization events [86, 4]. We considered each time-step in the simulation as an interval of size 1, and randomly positioned each speciation event in this interval (uniformly at random); this avoids multiple simultaneous speciation events.

In the three region model that includes a bridge region, we modeled the reduced prediction of a defined niche space in the bridge region (Fig. 2c) as a lack of region-dependent parameter rates. Therefore, the probabilities of *λ* and *µ* for genera occupying the bridge region were obtained from the region-dependent rates of the region from which the genera colonized the bridge.

#### 5.4.2 Simulation output and evaluation

We track the evolutionary history and geographic occurrence of all genera throughout the simulation. At the end of the simulation, we generate the phylogeny of the remaining genera using the tracked evolutionary history. We used the final distribution of genera to the different regions to compute SES-MPD values for each region. A null distribution was generated as described above in order to compute the standard effect size (SES) of MPD by permuting the assignment of each tips of the phylogeny to one of the tree regions (500 permutations were used).

#### 5.4.3 Running simulations

Simulations were ran for 5500 time steps. For model parameters, we selected *γ*, *µ* and *λ* values that generate a number of genera at the end of the simulation that is broadly comparable to the number of genera in our analysis of tree species (a few hundreds to 1–2 thousands) (Fig. S11). For speciation rates, we used three parameters: *λ* = 1*×*10*^−^*^3^, 5*×*10*^−^*^4^, and,1*×*10*^−^*^4^. For extinction rates, we used five parameters, defined as fractions of the speciation rates: *µ* = 0, 0.15, 0.3, 0.45 and 0.6; these values fall within the range of extinction rates relative to speciation rates previously reported for plant species [87, 88]. For colonization rates, we used four parameters: *γ* = 1 *×* 10*^−^*^4^, 1 *×* 10*^−^*^3^, 1 *×* 10*^−^*^2^ and 1 *×* 10*^−^*^1^; these rates span several orders of magnitudes, ranging as low as the speciation rates we used and as high as 0.1 colonization events per genera per time step. The high colonization rate values align with rates previously reported for tree species between tropical and subtropical regions [9]. The parameter choice was done to ensure a wide span of the parameter space with plausible parameter values, rather than restrict the space to realistic parameter combinations, because our goal with the model simulations was to understand the effect of the bridge region and evolutionary parameters on the emergence of latitudinal gradients. Taking all combinations of these parameter values, (two speciation parameters for the tropics and temperate regions, two extinction parameters for the tropics and temperate regions, and either four or two colonization parameters for the three- and two-region models) we have a parameter space of 57600 parameter configurations for the three-region model and 3600 configurations for the two-region model.

We retained only simulations that produced phylogenetic trees between 20 to 2000 surviv-ing genera (phylogenetic tips). For each parameter configuration, we generated 100 simula-tion replicates. We ran simulations until 100 successful simulations were obtained— defined as producing phylogenetic trees with 20 to 2000 surviving genera. Because of computational limitations, a maximum of 2000 simulations were allowed per parameter configuration, and if 100 “successful” simulations (with 20–2000 genera) were not achieved within this limit, this parameter configuration was excluded. As a result, we retained 97% of the parameter space for the two-region model and 87% for the three-region model.

## 6 Acknowledgments

This study was supported by the the JNF Climate Scholarship and the Minerva Center for the Study of Population Fragmentation. JTW was supported by an NSERC Discovery Grant.

## Supplementary Figures

**Figure S1:**
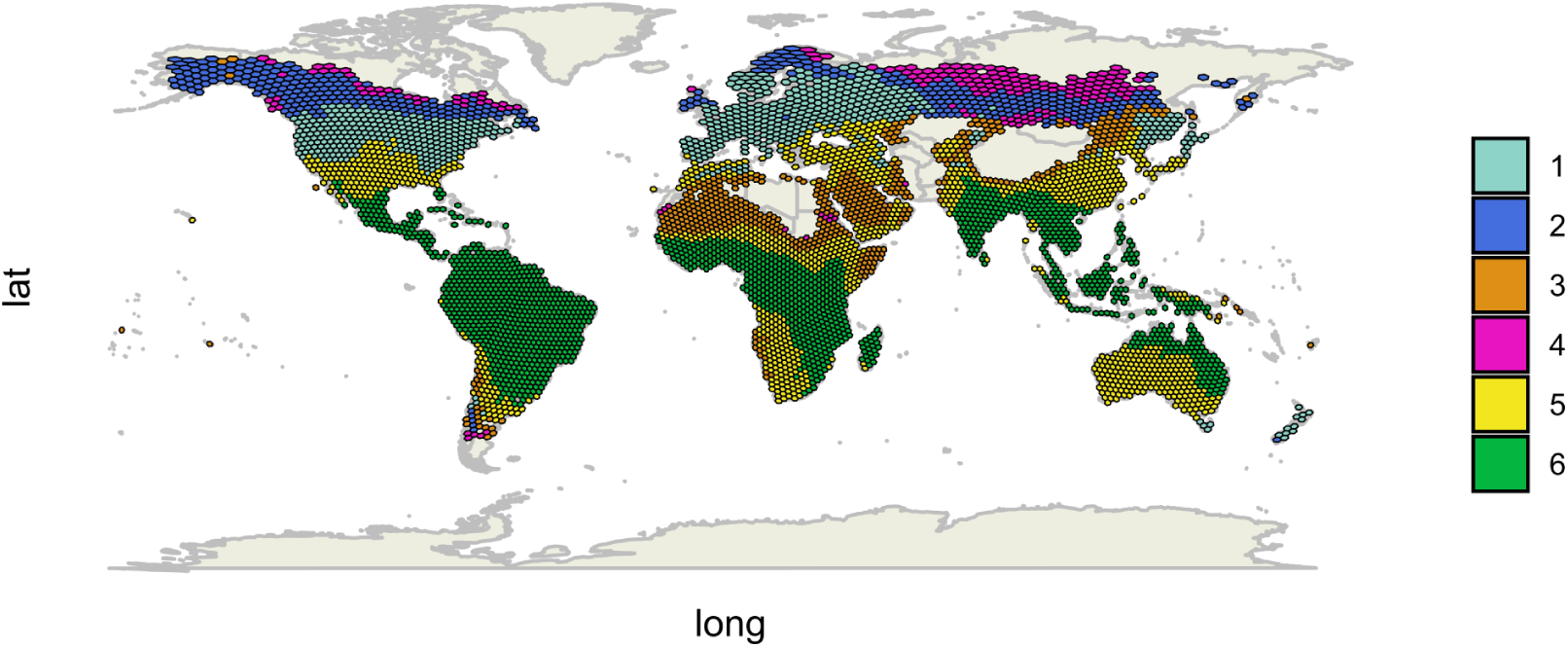
Phyloregion of tree species, with *k* = 6 clusters.

**Figure S2:**
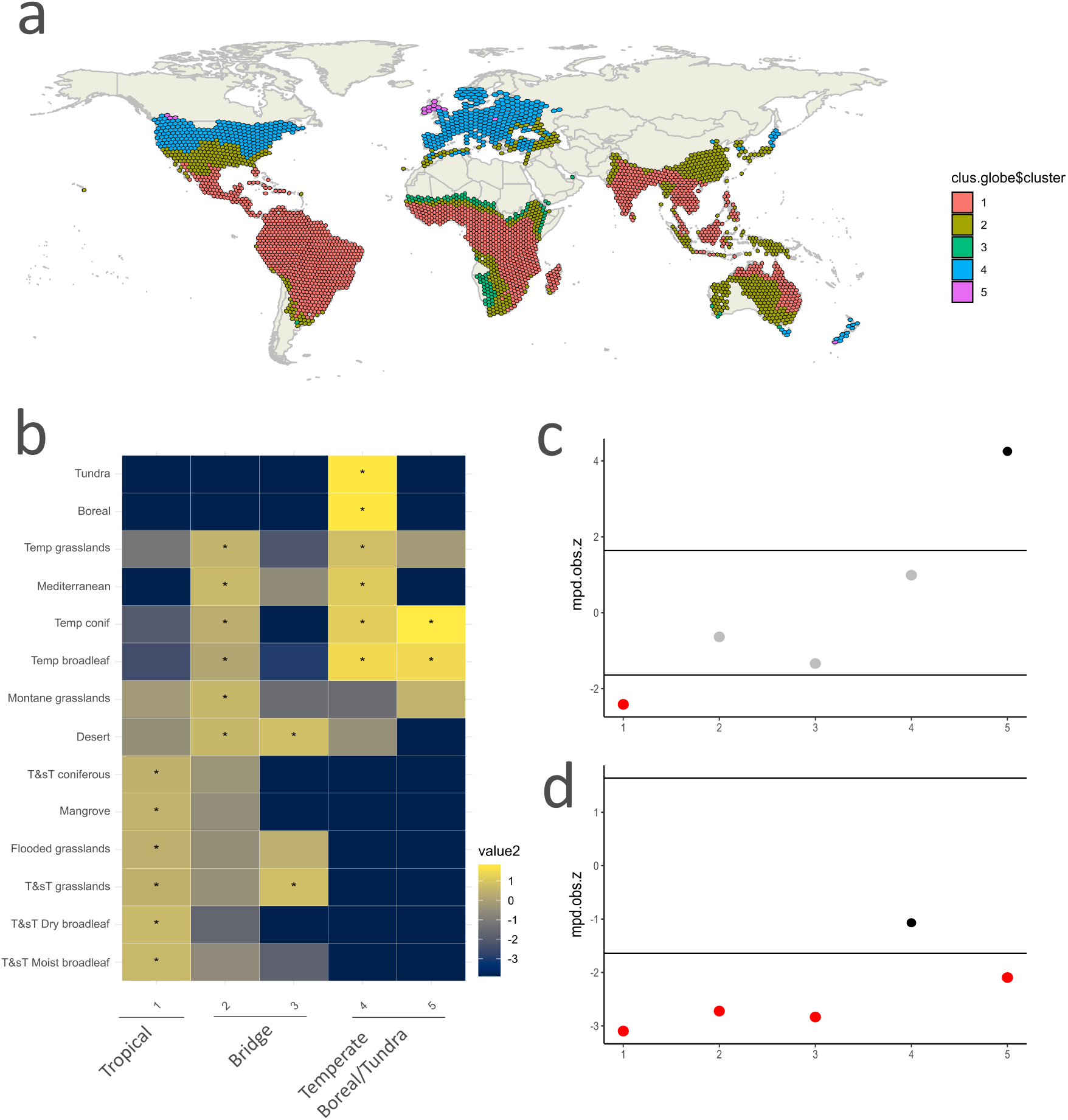
Generating phyloregions under a conserved filtering constrain. Generating phyloregion patterns of tree species, when accounting only for cells containing more than 30 species. (a) Phyloregion map distribution with *k* = 5 clusters (b) Proportion of overlap between each ecological biome and phyloregions, compared to a randomized phyloregion bootstrap. The heatmap shows the natural logarithm of the ratio between the actual overlap and the bootstrap overlap, with yellower colors signifying higher than random overlap. Statistical significance of high overlap is marked with an asterisk (*p <* 0.01). (c) Phylogenetic diversity of each of the phyloregions, measured as SES-MPD Z-scores. (d) Phylogenetic diversity of the phyloregions accounting only for angiosperms.

**Figure S3:**
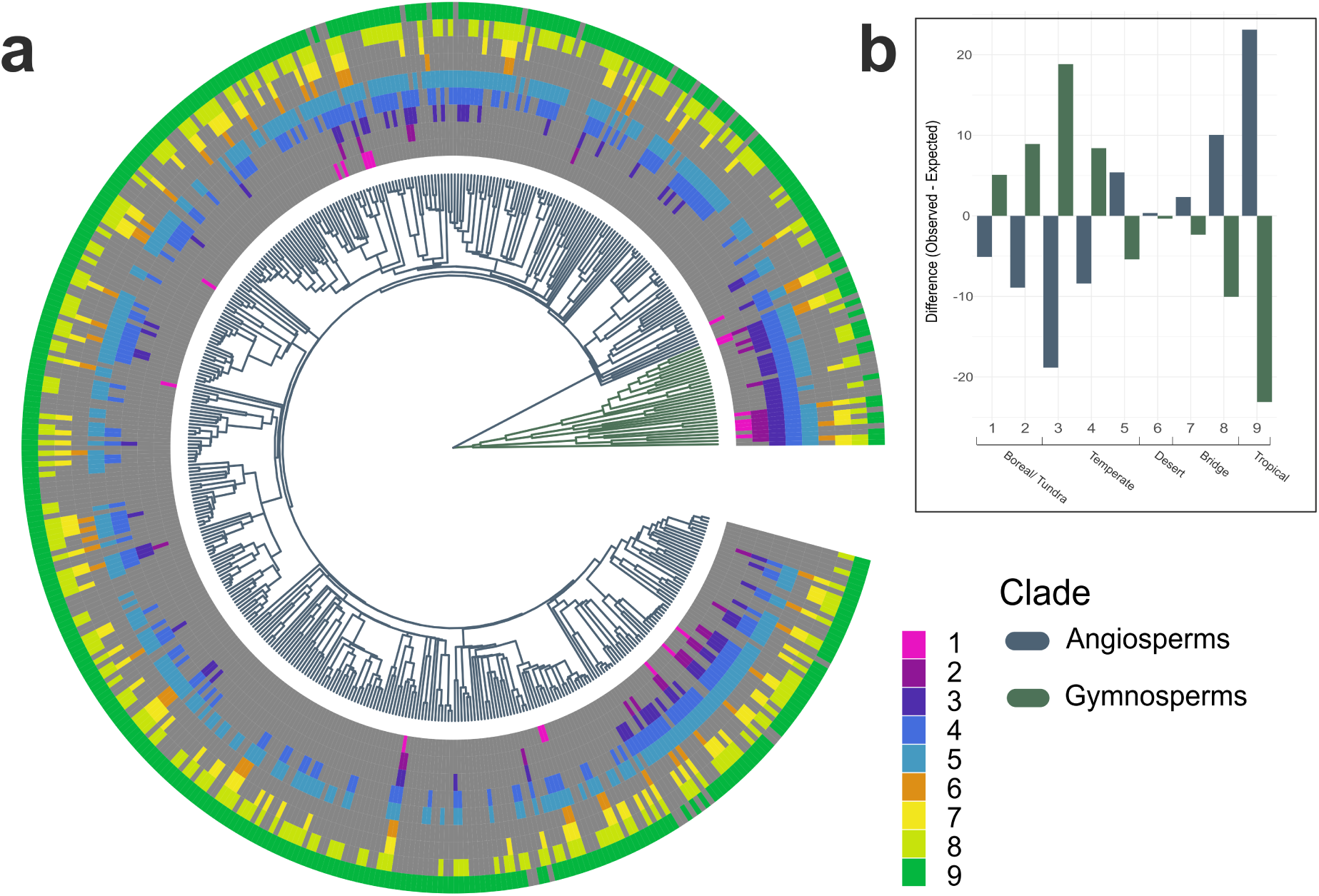
Phyloregion taxonomic composition. a) Phylogenetic tree of woody-species genera used to compute phyloregions, subdivided between gymosperms and angiosperms. Outer layers represent the presence or absence of each genus (phylogenetic tip) within each of the phyloregions. b) Difference between observed and expected presence of the taxonomic groups in each of the phyloregions, as a function of the total occupancy of genus/phyloregion.

**Figure S4:**
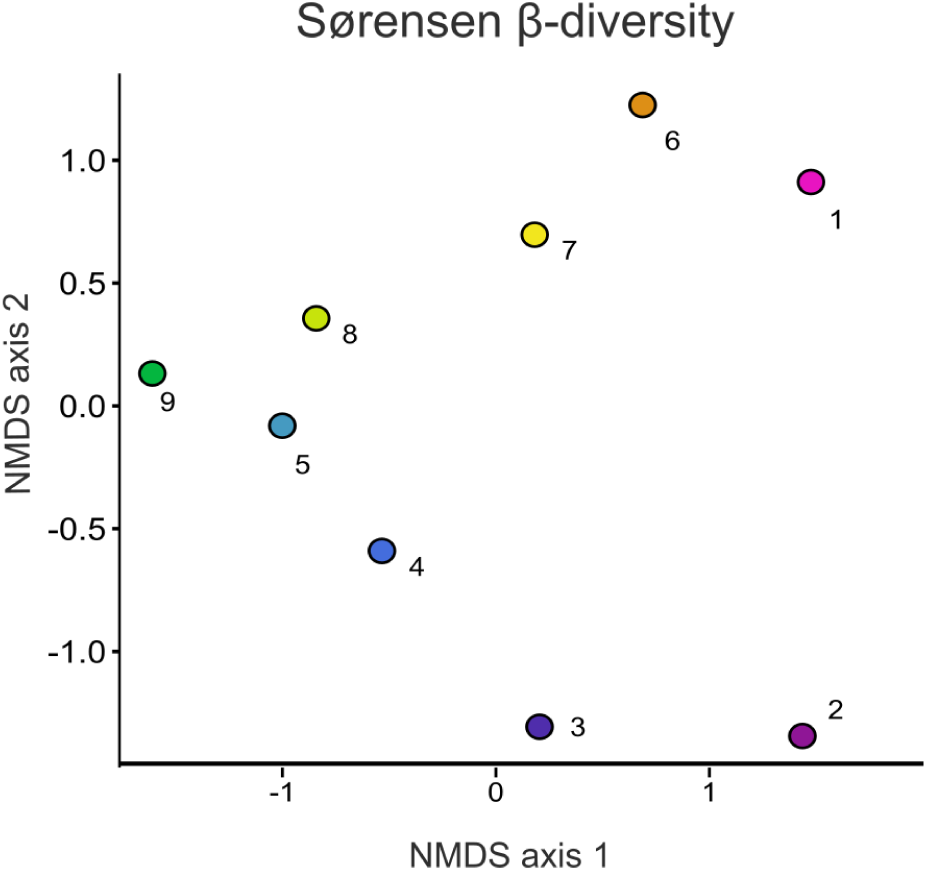
Community relationship amongst phyloregions. NMDS ordination plot of the distance matrix between phyloregions measured using Sørensen’s *β*-diversity index

**Figure S5:**
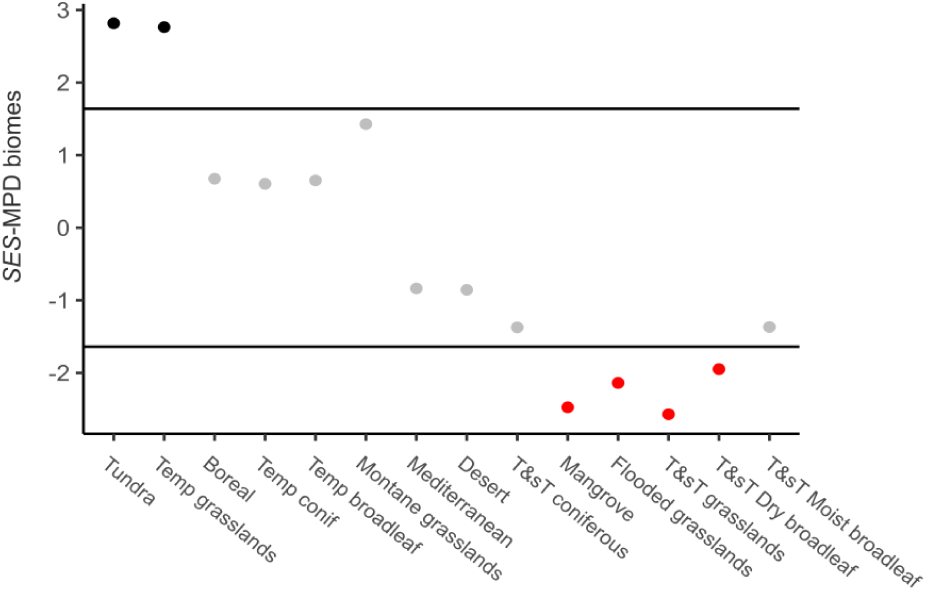
Phylogenetic diversity measured as *SES −MPD* along the biomes. Corresponding mean phylogenetic distance (SES-MPD, *Z-score*). Regions with SES-MPD *Z-*scores *< −*1.96 (red dots) have a significant phylogenetic clustering of tree species, phyloregions with SES-MPD *Z-*score *>* 1.96 (black dots) have significant phylogenetic over-dispersal of tree species.

**Figure S6:**
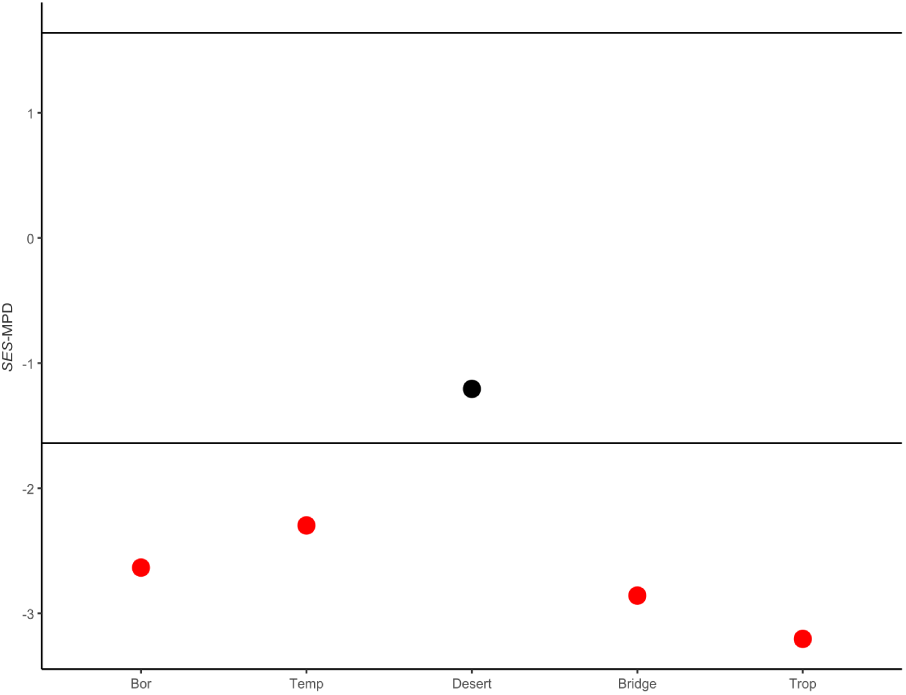
Phylogenetic diversity measured as *SES − MPD* when accounting only the distribution of angiosperms (excluding gymnosperms) Corresponding mean phylogenetic distance (SES-MPD, *Z-score*). Regions with SES-MPD *Z-*scores < −1.96 (red dots) have a significant phylogenetic clustering of tree species, phyloregions with SES-MPD *Z-*score *>* 1.96 (black dots) have significant phylogenetic over-dispersal of tree species.

**Figure S7:**
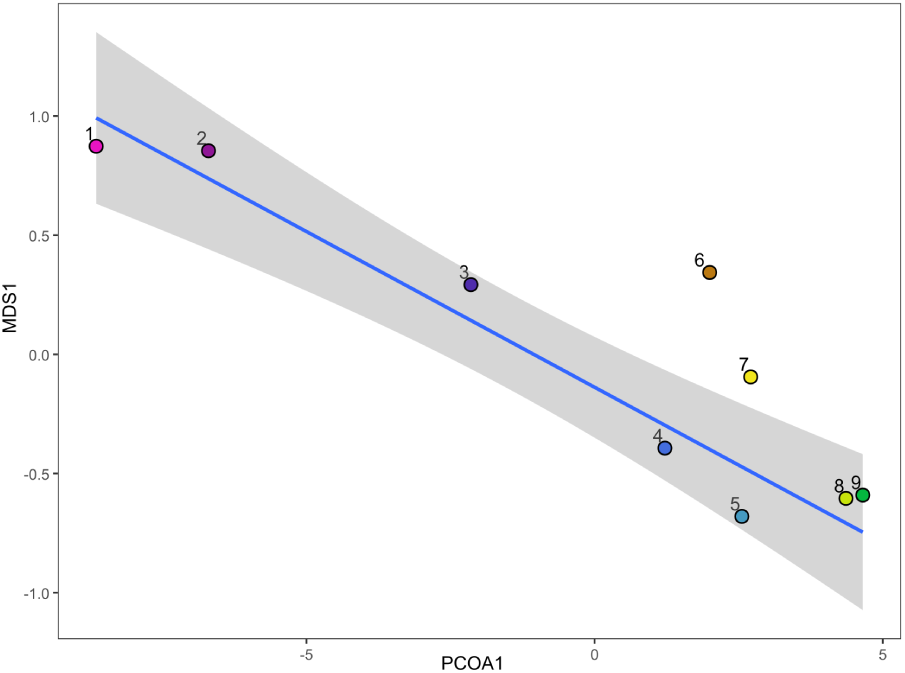
Correlation between the first axis of NMDS analysis of genera turnover (Supp. Figure S4) and PCoA analysis of phylogenetic turnover (Figure 2b)

**Figure S8:**
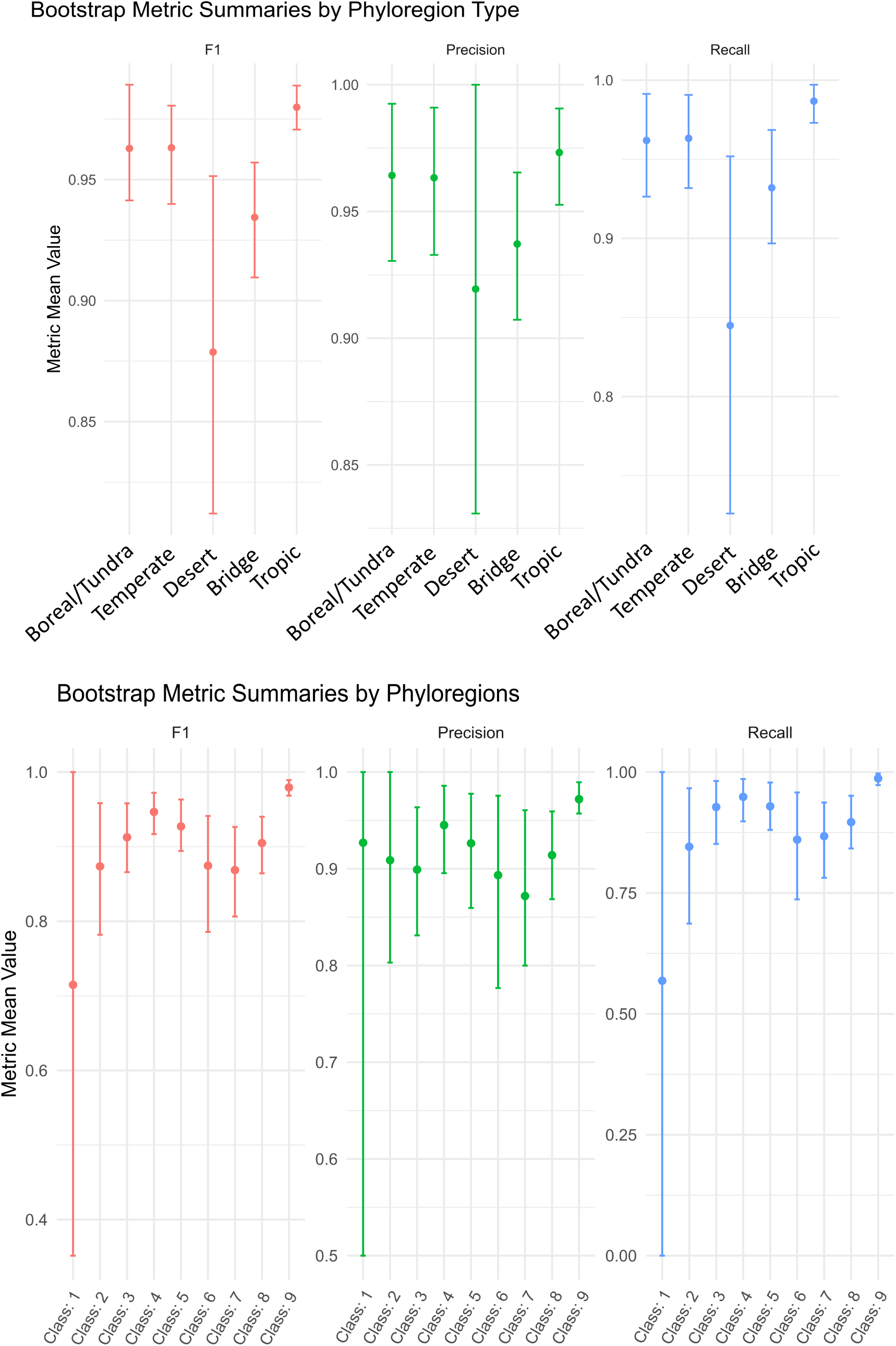
Phylogenetic and climatic niche conservatism. F-1, Precision and Recall rates for (a) each phyloregion and (b) the cluster of phyloregions from a random forest model.

**Figure S9:**
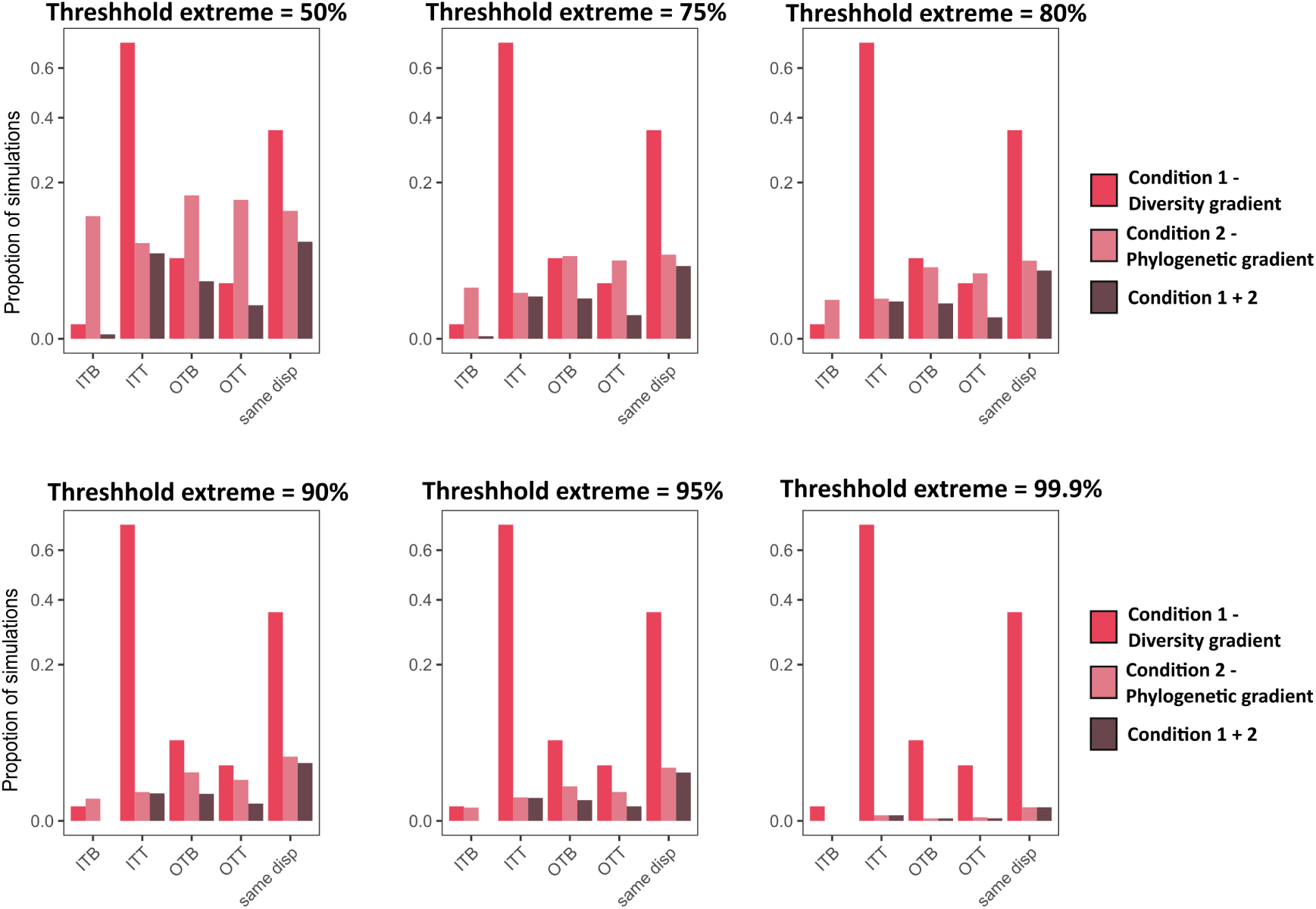
Scenarios in which latitudinal gradients emerge. The proportion of the simulations for which the latitude diversity gradient (Condition 1),the latitude phylogenetic gradient (Condition 2) and a combination of both emerged for each type of colonization dynamic under varying threshholds of MPD values, where the highest X% of tropical MPD values and the lowest X% of temperate MPD values were selected

**Figure S10:**
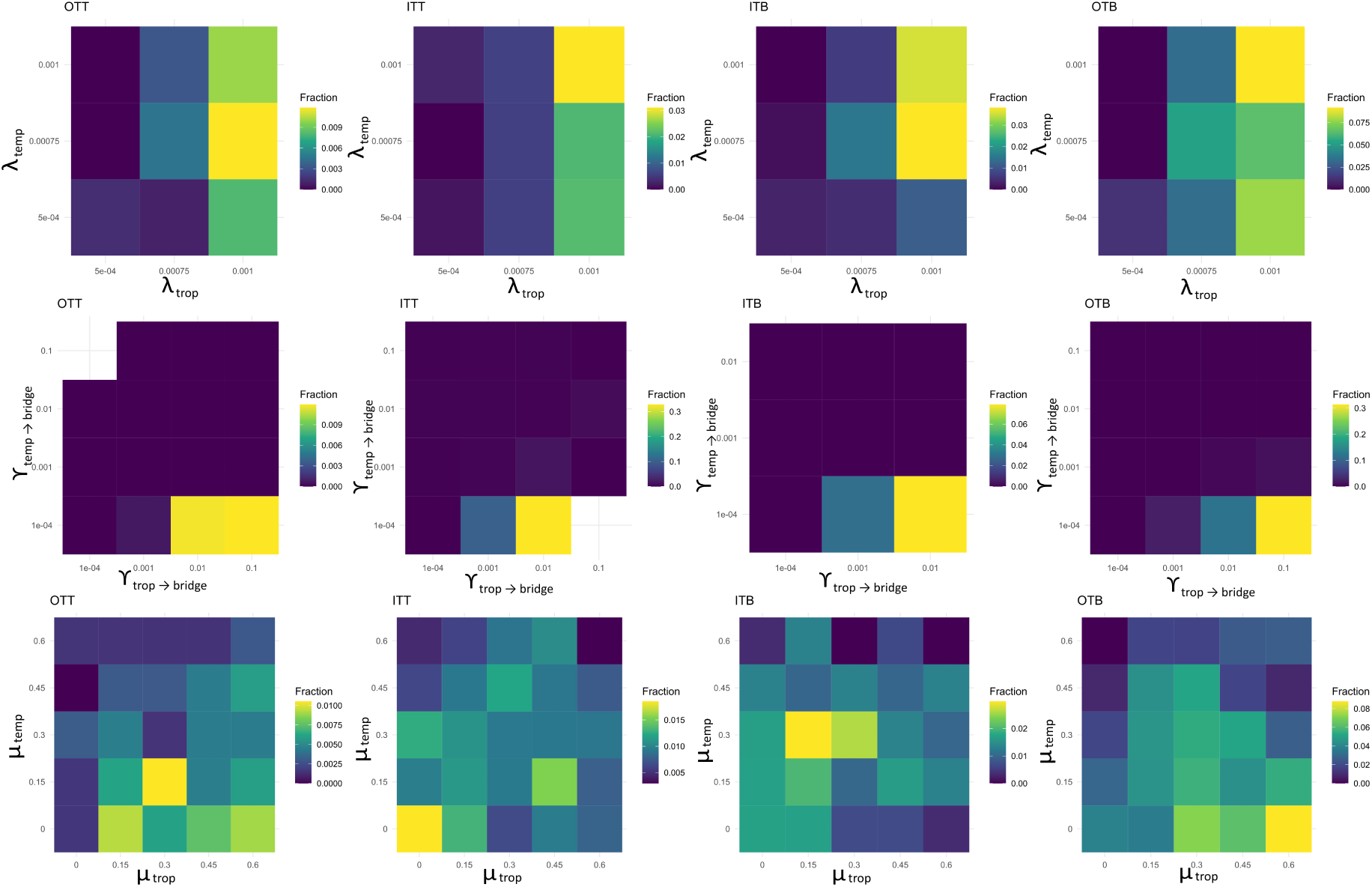
Proportion of simulations where a range of parameters enable the expected output for the latitude phylogenetic and diversity gradients to occur. The extinction rates (*µ*) shown are the proportions of speciation rates (*λ*)

**Figure S11:**
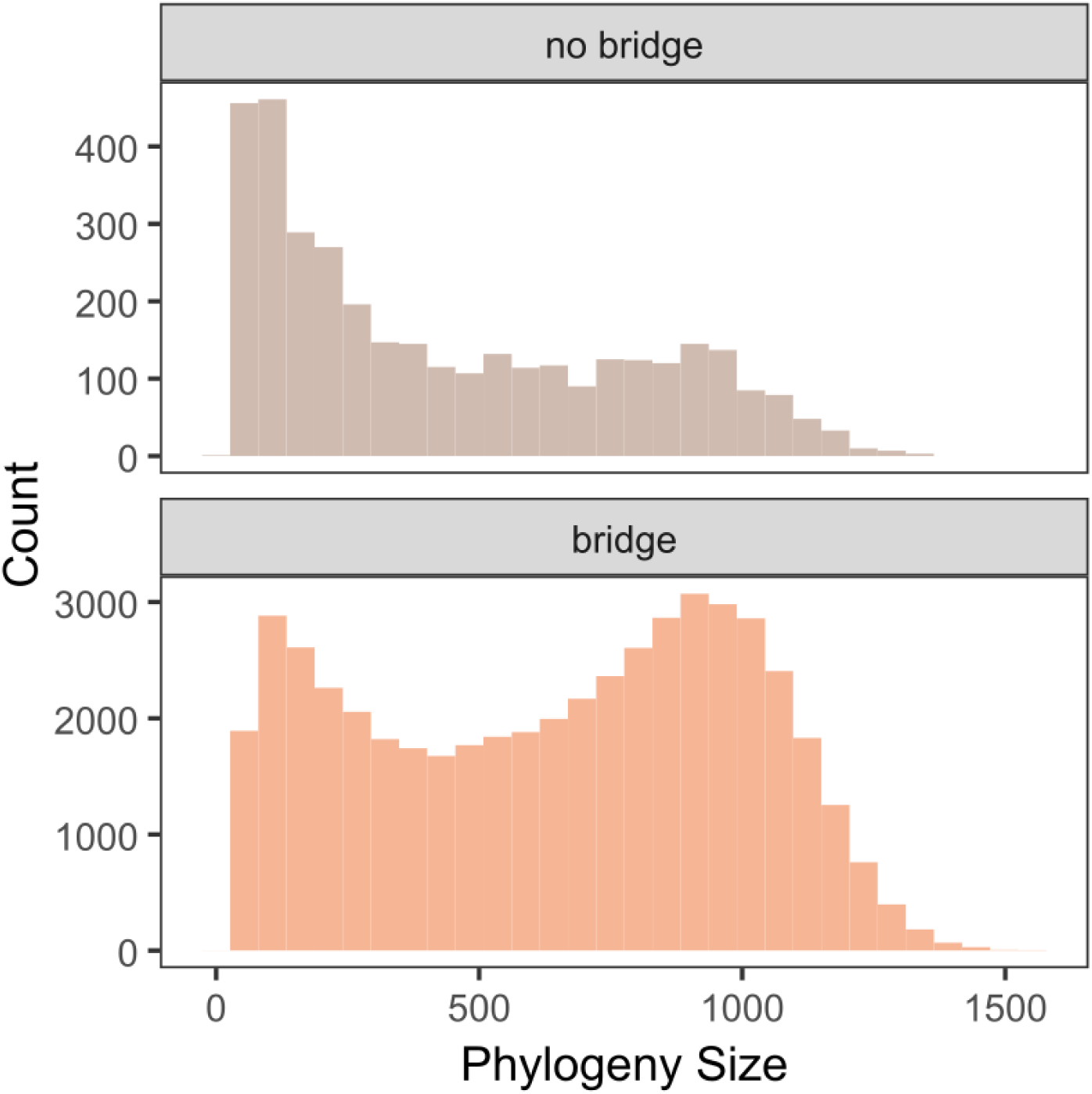
Histograms showing the sizes of the phylogenetic trees generated from the models with and without a bridge region.

